# Fructose 1,6-bisphosphate sensing by pyruvate kinase isozymes M2 (PKM2) controls MyoD stability and myogenic differentiation

**DOI:** 10.1101/2020.12.22.424062

**Authors:** Minchul Kim, Yao Zhang, Carmen Birchmeier

## Abstract

Glucose exerts beneficial effects on myogenesis and muscle physiology. However, the mechanisms by which glucose regulates myogenesis remain ill-defined or incompletely understood. Here, we show that low glycolysis destabilizes MyoD protein, a master myogenic transcription factor. Intriguingly, MyoD is not controlled by the cellular energy status per se, but by the level of fructose 1,6-bisphosphate, an intermediate metabolite of glycolysis. Fructose 1,6-bisphosphate is sensed by pyruvate kinase M2 (PKM2). In the presence of fructose 1,6-bisphosphate, PKM2 form tetramers that sequester the Huwe1 E3 ubiquitin ligase to the cytoplasm. Reduced fructose 1,6-bisphosphate levels dissociate the tetramer, releasing Huwe1 into the nucleus where it targets MyoD for degradation. Genetic or pharmacological modulation of PKM2-Huwe1 axis restores myogenic differentiation in glucose restricted conditions. Our results show that glucose metabolism directly regulates protein stability of a key myogenic factor and provide a rationale for enhancing myogenesis.

## Introduction

Glycolysis and oxidative phosphorylation are basic elements of cellular bioenergetics. However, a shutdown of glycolysis can severely affect cellular functions even in the presence of other fuels that produce ample energy. This is because intermediates of glycolysis have additional, non-bioenergetic roles. For instance, nuclear-cytoplasmic levels of the intermediate metabolite acetyl-CoA is rate-limiting for histone acetylation (Moussaieff et al., 2015). Thus, glycolytic flux and the resulting concentration of acetyl-CoA affects the epigenetic landscape, cell proliferation and differentiation. Additional glycolytic metabolites directly interact with or post-translationally modify proteins, thereby altering their activities (Ashizawa et al., 1991, Dayton et al., 2016, Bollong et al., 2018, Li et al., 2019). Thus, glucose metabolism plays a crucial role in many biological processes, acting through bioenergetic and non-bioenergetic mechanisms to control tumorigenesis, tissue repair and the function of the immune system (Pearce and Pearce, 2013, Hsu and Sabatini, 2008, Cairns et al., 2011, Folmes et al., 2012).

The skeletal muscle is the major sink for blood glucose and an important tissue for glucose homeostasis. Muscle fiber growth can occur by accretion of new nuclei to the fiber, i.e. fusion of myoblasts derived from muscle stem cells, or by increased fiber protein synthesis. Glucose can affect both aspects, but the precise effect and associated mechanism are different at distinct stages of myogenesis.

Myogenic cells progress through several stages in order to differentiate that are characterized by the expression of distinct transcription factors (Yin et al., 2013, Brack and Rando, 2012, Comai and Tajbakhsh, 2014, Buckingham and Rigby, 2014). Adult muscle stem cells express the transcription factor Pax7, which is essential for the survival and maintenance of muscle stem cell pool (von Maltzahn et al., 2013, Oustanina et al., 2004). Upon activation, these cells start to express the transcription factors MyoD and/or Myf5. Activated stem cells can either return to quiescence or further progress into differentiation (Bentzinger et al., 2012). Finally, myoblasts turn on the transcription factor myogenin (MyoG), a marker of terminal differentiation, and exit the cell cycle to fuse into myotubes. Previous studies have shown that the glucose promotes activation of quiescent muscle stem cells and differentiation of myoblasts in part by regulating histone acetylation (Fulco et al., 2008, Yucel et al., 2019, Ahsan et al., 2020, Theret et al., 2017). In differentiated myotubes, glucose inhibits the Foxo transcription factors whose activity induces the expression of muscle-specific ubiquitin ligases that result in atrophy (Sandri et al., 2004, Meng et al., 2017). Knowledge about the molecular mechanisms underpinning these effects can help to design better strategies to enhance myogenesis, for instance in a disease setting.

Here, we aimed to identify a direct mechanistic link between glucose and key myogenic factor(s) in order to better understand the molecular mechanism by which glucose promotes differentiation of myoblasts. We observe that the protein stability of MyoD depends on glucose and define here the responsible pathway. In particular, we show that fructose 1,6-bisphosphate inhibits the E3 ubiquitin ligase Huwe1 to stabilize MyoD. Fructose 1,6-bisphosphate is sensed by the M2 isoform of pyruvate kinase that in turn interacts with Huwe1 to control its subcellular location.

## Results and discussion

### Glucose regulates MyoD protein stability

Consistent with the previous report (Fulco et al., 2008), lowering of the glucose concentration in the medium to 5mM impaired differentiation of cultured C2C12 myoblast cells (Fig. 1A). This was accompanied by a reduced expression of the terminal differentiation markers troponin-T and muscle creatine kinase (Mck). Interestingly, MyoD protein levels were drastically reduced in low glucose (Fig. 1A), but MyoD mRNA was unchanged (Fig. 1B). In contrast, both MyoG protein and mRNA were significantly downregulated. MyoG is a target of MyoD (Deato et al., 2008), indicating that the strong downregulation of MyoD protein is an important mechanism by which low glucose impairs differentiation.

**Figure 1.**
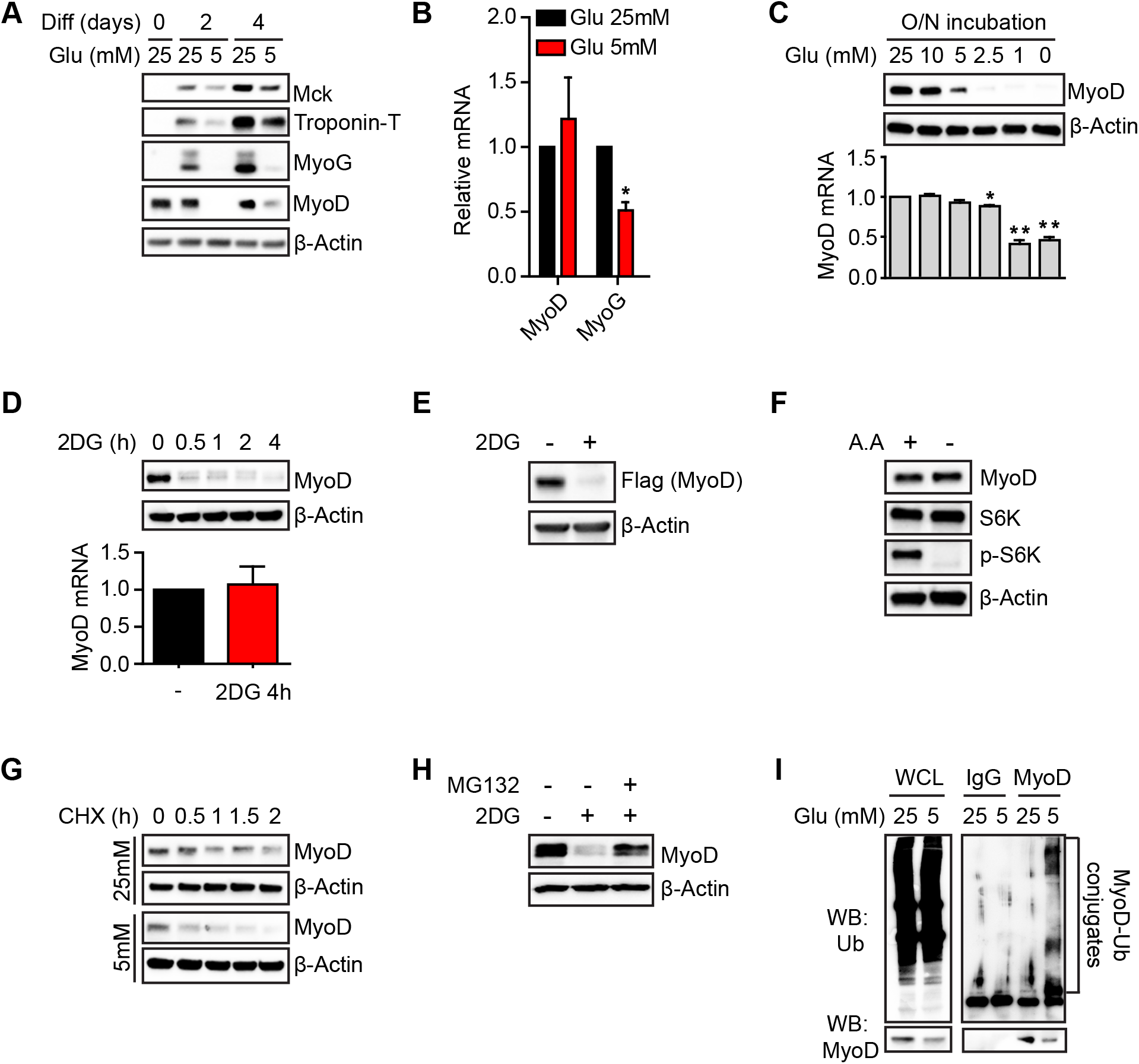
Glucose regulates MyoD protein stability via ubiquitination. (A) C2C12 cells were differentiated at indicated glucose concentrations and analyzed by Western blotting. (B) Samples from (A) at day 2 were analyzed by RT-qPCR for MyoD or MyoG (n=3). (C) Cells were incubated in proliferation media (10% FBS) with a range of glucose concentrations overnight and analyzed by Western blotting or RT-qPCR (n=3). (D) Cells were treated with 25mM 2DG in glucose free media for indicated time and analyzed by Western blotting or RT-qPCR (n=3). (E) Cells were transfected with Flag-MyoD construct and treated with 2DG for 4 hours before harvesting. (F) Cells were incubated in amino acid free media for one hour. Loss of p-S6K was used to confirm amino acid starvation response. (G) Cells were cultured in 25mM or 5mM glucose media for one day and chased after 20μM CHX treatment. (H) Cells were treated with 2DG in the presence or absence of 20μM MG132 for 4 hours. (I) Cells were cultured in 25mM or 5mM glucose media for one day and incubated with 20μM MG132 for 4 hours. Cell lysates were immunoprecipiated with an anti-MyoD antibody to access ubiquitination. Western blot results were validated in two or more independent experiments. Error bars indicate S.E.M. Paired Student’s t-test (two-tailed) was performed. *, p<0.05. **, p<0.01.

To further investigate the regulation of MyoD by glucose, we cultured proliferating C2C12 cells in media with different glucose concentrations overnight. MyoD protein levels were similar in 25mM and 10mM glucose but were markedly reduced in 5mM and 2.5mM (Fig. 1C, western blot). In contrast, MyoD mRNA remained comparable within a range of 2.5-25mM glucose and was only affected in a pronounced manner when glucose levels decreased even further (Fig. 1C, graph). Thus, MyoD protein quantity is dynamically regulated in a range of glucose concentrations that is physiologically relevant (4-7mM in normally fed mice). MyoD protein was depleted within 4 hours when glycolysis was acutely blocked by 25mM 2-Deoxyglucose (2DG) in glucose-free medium, but MyoD mRNA was unaffected in this time window (Fig. 1D). Further supporting a non-transcriptional control, 2DG also depleted Flag-MyoD protein that was produced from a transfected plasmid (Fig. 1E). Unlike low glucose concentrations, amino acid starvation did not affect MyoD protein levels, indicating that a shutdown of glycolysis specifically regulates MyoD levels (Fig. 1F).

Since transcriptional mechanism cannot account for the changes in MyoD levels, we tested whether the stability of MyoD protein is affected by low glucose. Addition of cycloheximide, an inhibitor of translation, showed that the turnover of MyoD protein was higher in 5mM glucose than in 25mM (Fig. 1G). Furthermore, in the presence of the proteasome inhibitor MG132, MyoD protein levels were restored when glycolysis was acutely blocked by 2DG (Fig. 1H). Finally, we found that MyoD ubiquitin levels were substantially higher in cells cultured in 5mM glucose than in 25mM (Fig. 1I). We conclude that glucose regulates MyoD protein stability through the ubiquitination-proteasome pathway.

### Cellular energy stress does not regulate MyoD stability

We next sought to characterize the mechanism that mediates the effect of glucose on MyoD stability. AMP-activated kinase (AMPK) and Sirt1 (NAD^+^-dependent deacetylase) are well-established sensors of cellular energy stress, and account for many aspects of the biological responses to energy starvation (Canto et al., 2010, Fulco and Sartorelli, 2008). We mutated AMPKα1/2, the genes encoding the catalytic subunits of the AMPK complex using CRISPR-Cas9 to generate two independent mutant clones. MyoD was unstable in the presence of 2DG in both mutant clones (Fig. 2A). Furthermore, the specific Sirt1 inhibitor (EX527) did not restore MyoD protein stability (Fig. 2B). Thus, none of the well-known cellular energy sensors is responsible for the de-stabilization of MyoD when glycolysis is blocked.

**Figure 2.**
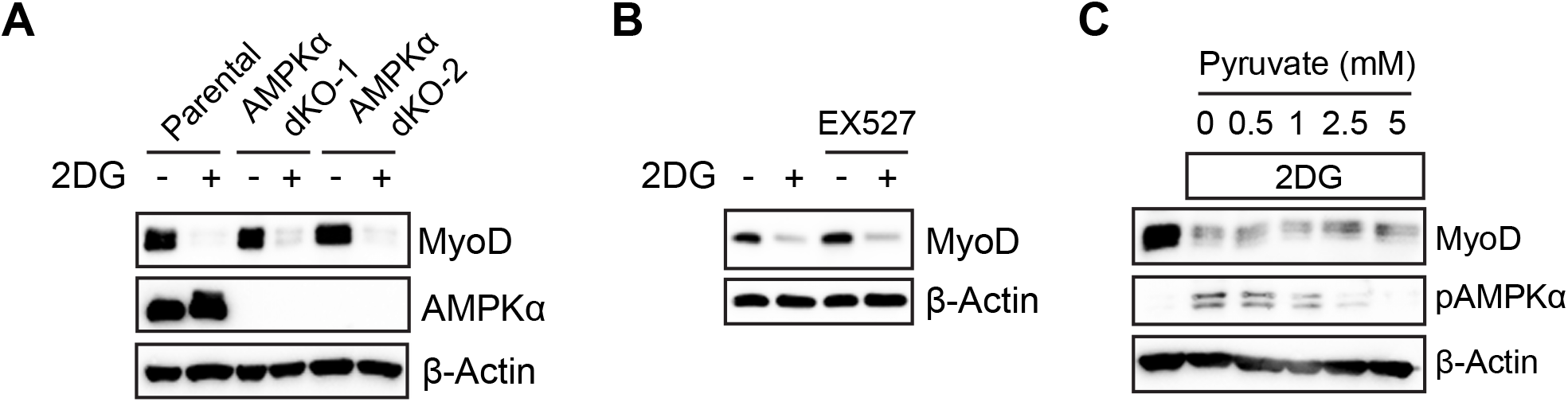
Cellular energy per se do not regulate MyoD. (A) Parental C2C12 or two Ampkα1/2 dKO clones were treated with 2DG for 4 hours. (B) Cells were treated with 2DG in the presence or absence of 10μM EX527 (Sirt1 inhibitor). (C) Cells were co-treated with increasing concentrations of pyruvate and 2DG for 4 hours. All results were validated in two or more independent experiments.

This raised the possibility that not energy stress *per se*, but the absence of glycolysis and thus the depletion of specific glycolytic intermediates regulates MyoD stability. To test this, we replaced glucose with pyruvate as energy source and analyzed phosphorylation of AMPK to monitor the cellular energy status. In the presence of 2DG, AMPK is phosphorylated, indicating that the cells experience energy stress. With increasing concentrations of added pyruvate, AMPK phosphorylation declined. In particular, at a pyruvate concentration of 2.5mM, AMPK phosphorylation had returned to baseline levels, indicating that the energy metabolism of the cell was restored. However, MyoD protein remained destabilized at all pyruvate concentrations (Fig. 2C). This result suggested that an intermediate of the glycolytic pathway produced by glucose but not pyruvate regulates MyoD stability (Fig. 3A).

**Figure 3.**
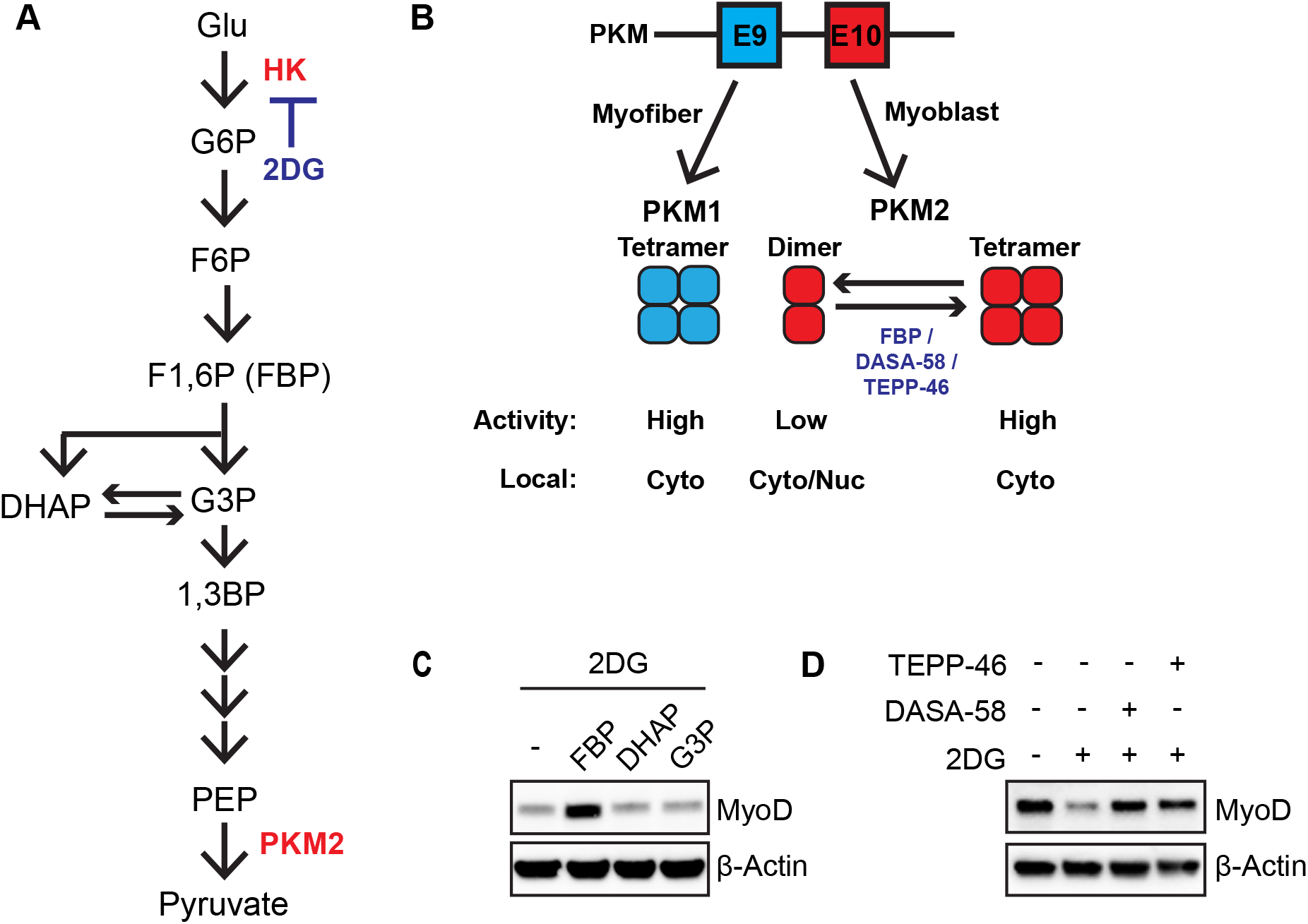
PKM2 is the responsible sensor of FBP for regulation of MyoD. (A) Diagram of glycolysis pathway. (B) Diagram illustrating the regulation of Pyruvate Kinase M gene by alternative splicing in myofibers and myoblasts, and different biochemical properties of PKM1 and PKM2. Myofibers use exon 9 to express PKM1 whereas myoblasts use exon 10 to express PKM2. FBP (fructose 1,6-bisphosphate), DASA-58 and TEPP-46 stabilize PKM2 tetramer. (C) Cells were permeabilized with Streptolysin-O (SLO), and treated with 2DG and indicated metabolites (200μM) for 2 hours. DHAP, dihydroxyacetone phosphate. G3P, glyceraldehyde 3-phosphate. (D) Cells were co-treated with 2DG and PKM2 tetramer stabilizing reagents (100μM) for 4 hours. All results were validated in two or more independent experiments.

### Fructose 1,6-bisphosphate sensing by PKM2 stabilizes MyoD

Fructose 1,6-bisphosphate is produced from fructose-6-phosphate (F6P) at the third step of glycolysis. The concentration of fructose 1,6-bisphosphate can be sensed by its binding to the isoform 2 of the muscle Pyruvate Kinase (PKM2) (Fig. 3A). PKM2 is expressed in proliferating myoblasts, whereas due to alternative splicing the PKM1 isoform is produced in differentiated myofibers (David et al., 2010). Binding of fructose 1,6-bisphosphate to PKM2 induces tetramerization of the dimeric/monomeric PKM2, which augments its enzymatic activity and promotes its cytoplasmic localization. On the other hand, absence of fructose 1,6-bisphosphate dissociates the tetramer, resulting in lower enzymatic activity of PKM2 and its translocation into the nucleus (Dayton et al., 2016) (Fig. 3B). This delicate regulation of PKM2 activity and localization is known to be required for cell proliferation (Israelsen et al., 2013, Dayton et al., 2016). However, whether PKM2 integrates cellular metabolic signals to regulate differentiation is unknown.

Interestingly, supplying fructose 1,6-bisphosphate, but not its downstream products G3P (glyceraldehyde 3-phosphate) or DHAP (dihydroxyacetone phosphate), protected MyoD from degradation in the presence of 2DG (Fig. 3C). To test if dissociation of the PKM2 tetramer contributes to the regulation of MyoD turnover, we used DASA-58 or TEPP-46, two compounds that stabilize the PKM2 tetramer (Anastasiou et al., 2012). Remarkably, both DASA-58 and TEPP-46 stabilized MyoD in the presence of 2DG (Fig. 3D). Thus, in the metabolic pathway that regulates MyoD stability, fructose 1,6-bisphosphate has a key regulatory role and is sensed by PKM2.

### The E3 ubiquitin ligase Huwe1 regulates MyoD stability during glucose deprivation

To gain further mechanistic insight, we next sought for the responsible ubiquitin E3 ligase that destabilizes MyoD in the presence of low glucose. Several E3 ligases including Huwe1 have been suggested to target MyoD (Noy et al., 2012). When Huwe1 was depleted using two different siRNAs, MyoD protein remained stable in the presence of 2DG (Fig. 4A). In contrast, two other E3 ligases known to target MyoD, Atrogin-1 or Upf1 (Lagirand-Cantaloube et al., 2009, Feng et al., 2017), were not responsible for MyoD degradation in response to 2DG: inhibition of Atrogin-1 activity by MLN4924 or depletion of Upf1 by siRNA failed to protect MyoD from degradation in response to 2DG in myoblasts (Fig. EV1A). Thus, Huwe1 destabilizes MyoD in low glucose.

**Figure 4.**
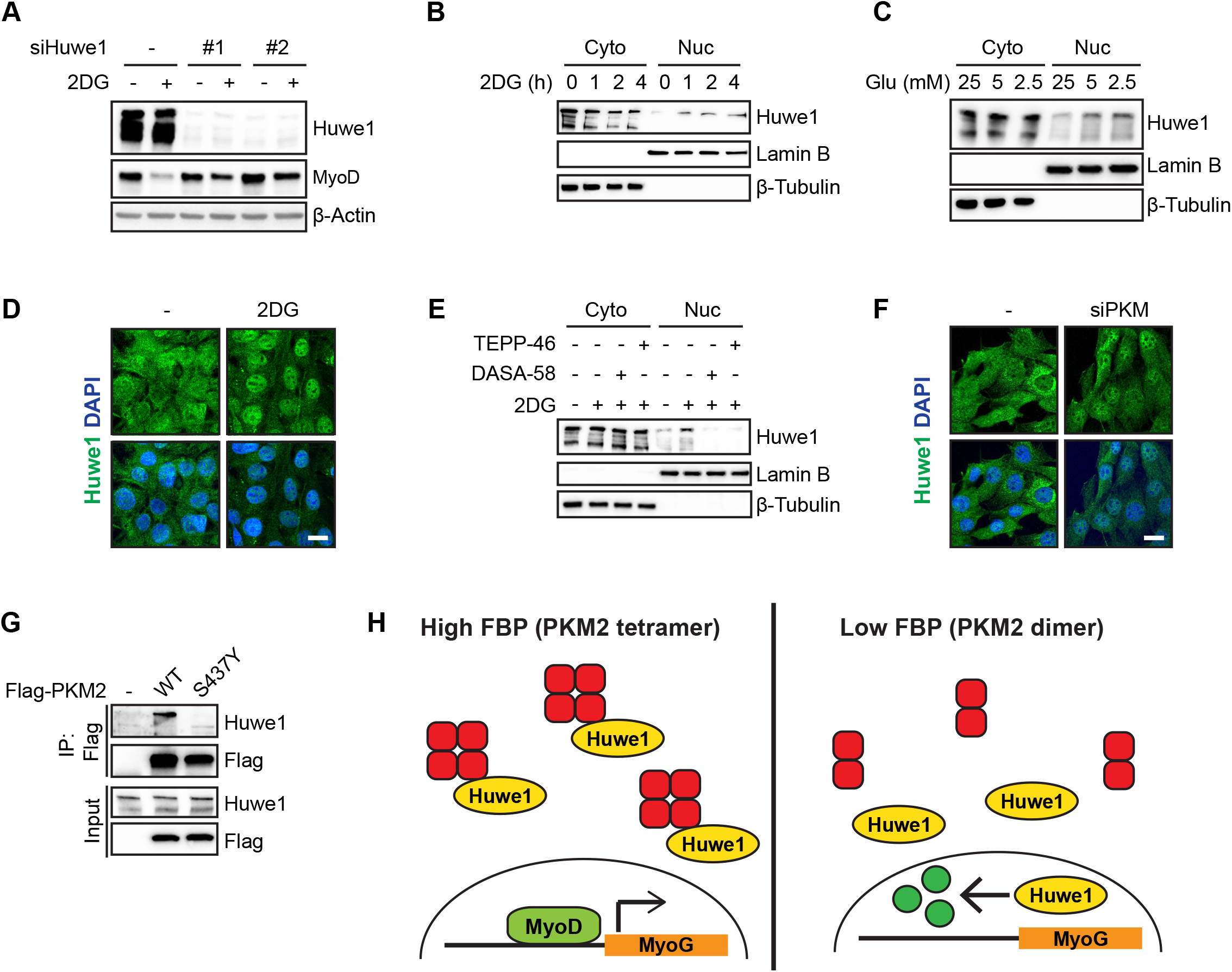
Huwe1 E3 ubiquitin ligase translocates to the nucleus following glycolysis inhibition, which is opposed by PKM2 tetramer. (A) C2C12 cells were transfected with two different siRNAs against Huwe1 and treated with 2DG for 4 hours. (B) Cells treated with 2DG for indicated time were fractionated to cytosolic and nuclear lysates. Lamin B is a nuclear marker and β-Tubulin is a cytosolic marker. (C) Cells were incubated in proliferation media with different glucose concentrations for one day and fractionated as in (B). (D) After 4 hours treatment with 2DG, cells were immunostained for Huwe1. Scale bar, 20μm. (E) Cells were treated as indicated for 4 hours and fractionated. (F) Control or PKM2 siRNA transfected cells were immunostained for Huwe1. Scale bar, 10μm. (G) Endogenous PKM2 was depleted using PKM2 siRNA (#1 in Fig. EV2C) followed by transfection with Flag-tagged human PKM2 WT or PKM2 S437Y mutant. Lysates were immunoprecipitated using antibody against Flag. (H) Model of MyoD regulation by FBP-PKM2-Huwe1 axis. FBP, fructose 1,6-bisphosphate. All results were validated in two or more independent experiments. Scales in (B) and (F), 10μm.

### PKM2 tetramer regulates the nucleocytoplasmic localization of Huwe1

Huwe1 is normally located in the cytoplasm, whereas MyoD is mostly nuclear. Nuclear translocation of Huwe1 has been reported in response to DNA damage or during spermatogenesis (Wang et al., 2014, Bose et al., 2017). Interestingly, Western blotting indicated that the levels of nuclear Huwe1 increased when cells were either treated with 2DG (Fig. 4B) or cultured in low glucose (Fig. 4C). We confirmed this by immunostaining experiments. In untreated cells, Huwe1 showed a heterogeneous distribution: in most cells Huwe1 displayed a diffuse cytoplasmic location, but occasionally cells with nuclear Huwe1 could be observed. In contrast, we observed a pronounced enrichment of Huwe1 in nuclei of 2DG treated cells (Fig. 4D). Critically, DASA-58 or TEPP-46 blocked nuclear Huwe1 translocation in response to 2DG (Fig. 4E), raising the possibility that the PKM2 tetramer binds Huwe1 and thus sequesters it in the cytoplasm.

We investigated this further by depleting PKM2 using siRNA, which resulted in an enhanced nuclear localization of Huwe1 (Fig. 4F). Immunostaining in Huwe1 siRNA transfected cells confirmed the specificity of Huwe1 antibody (Fig. EV2A). The increased level of nuclear Huwe1 was accompanied by a decreased basal level of MyoD in the absence of 2DG (Fig. EV2B). Notably, the degree of reduction of MyoD levels in the absence of 2DG correlated with the efficiency of the siRNA-induced PKM2 downregulation, indicating that incomplete depletion of PKM2 might have resulted in the presence of residual MyoD. In accordance, remaining MyoD was still destabilized by 2DG. PKM2 depletion did not affect basal MyoD mRNA levels (Fig. EV2C). Of note, despite multiple attempts using a number of gRNA sequences we failed to obtain PKM2 knock-out clones, retrieving only clones with reduced PKM2 protein levels, possibly heterozygous mutant clones. Therefore, PKM2 appears to be essential in C2C12 cells.

Next, we assessed whether PKM2 and Huwe1 physically interact. We found that Flag-tagged wildtype PKM2 co-immunoprecipitated endogenous Huwe1. In contrast, the S473Y mutant of PKM2 that is unable to bind fructose 1,6-bisphosphate (Chaneton et al., 2012) failed to interact with Huwe1 (Fig. 4G). Whether PKM2 and Huwe1 interact directly or via other protein(s) needs further investigation. Taken together, these results suggest that in the presence of fructose 1,6-bisphosphate, the PKM2 tetramer acts as a sink of Huwe1 in the cytoplasm (see a summary of the model in Fig. 4H). When cellular fructose 1,6-bisphosphate drop, the PKM2 tetramers dissociate and release Huwe1, which unleashes nuclear Huwe1 activity.

### Depletion of Huwe1 restores myogenic differentiation in low glucose condition

We next tested if inhibition of the PKM2-Huwe1 pathway reinstates differentiation in glucose restricted conditions. Depletion of Huwe1 by siRNA partially rescued myogenic differentiation in low glucose as assessed by MyoG immunostaining, a marker for onset of differentiation (Fig. 5A; quantified in 5B). In contrast, differentiation in high glucose was unaffected when Huwe1 was transiently depleted by siRNA. This is in contrast to a previous study using stable Huwe1 depletion by shRNA, which had reported impaired differentiation of C2C12 cells (Li et al., 2015). We also observed attenuated differentiation when we used shRNA targeting the same sequence of the Huwe1 transcript that is recognized by the siRNA used (Fig. EV3), indicating that long-term inhibition of Huwe1 has indirect effects on myogenic differentiation. Consistent with a previous report, PKM2 activators (DASA-58 or TEPP-46) strongly impaired cell proliferation when used on sub-confluent cells (Anastasiou et al., 2012). We therefore used confluent cultures and induced differentiation with or without TEPP-46 in the presence of low/high glucose. The addition of TEPP-46 partially restored myogenic differentiation in low glucose (Fig. 5A and 5C). In summary, both Huwe1 siRNA and PKM2 activators substantially but not completely restored MyoG induction in low glucose. The remaining effect might be mediated by other pathways such as AMPK and/or histone acetylation.

**Figure 5.**
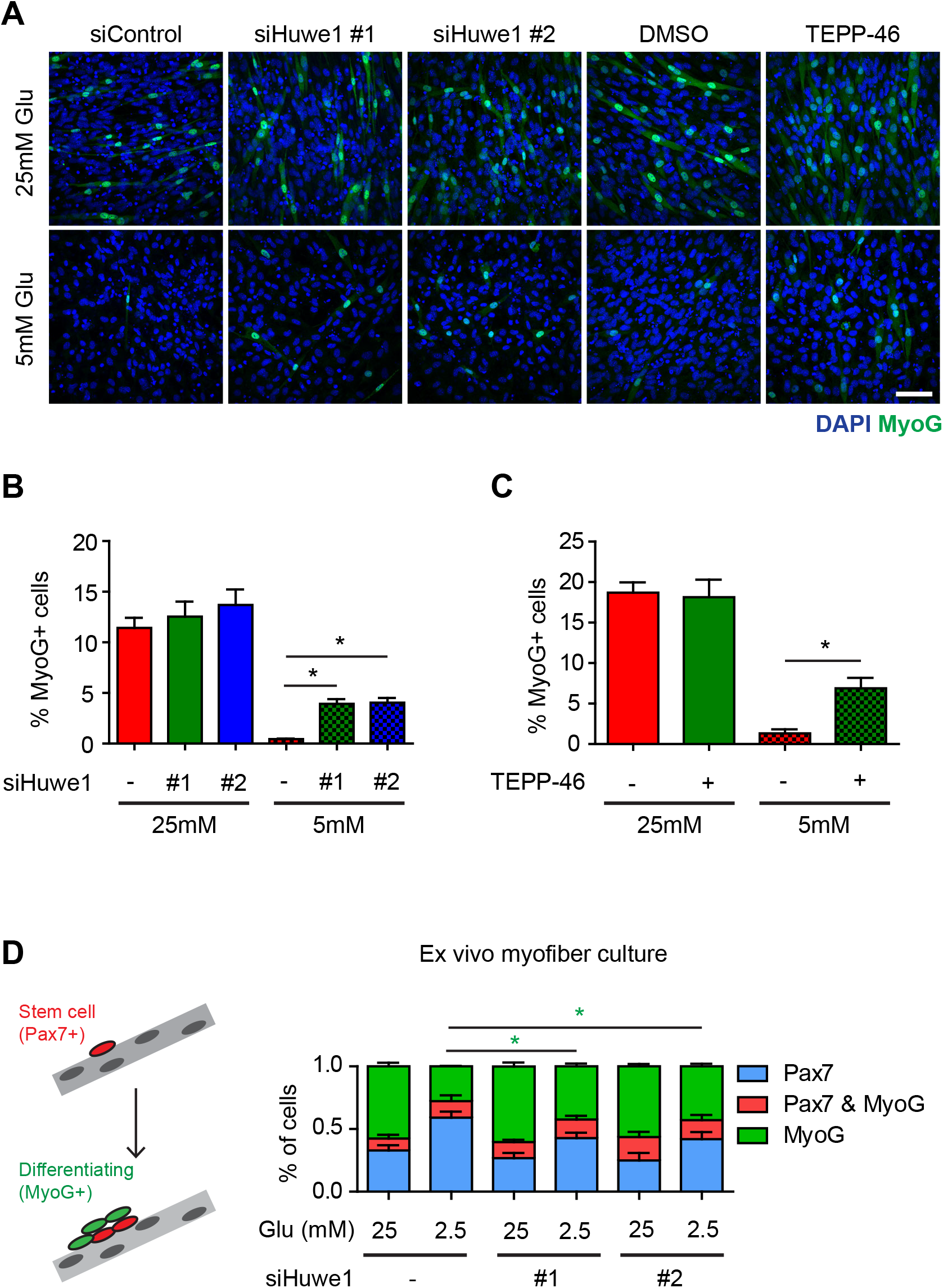
Inhibition of PKM2-Huwe1 axis restores myogenic differentiation in glucose restricted conditions. (A) Representative images of MyoG immunostaining in (B) and (C). Scale bar, 20μm (B) Huwe1 was depleted by siRNA in C2C12 cells, and differentiation was induced in 25mM or 5mM glucose media. Percentages of MyoG positive cells were quantified over at least 500 nuclei per sample (n=3). (C) Confluent C2C12 cells were induced to differentiate in 25mM or 5mM glucose media with or without 5μM TEPP-46. Percentages of MyoG positive cells were quantified as in (A) (n=3). (D) Left, diagram describing stem cell activation and differentiation in cultured myofibers. Right, isolated mouse EDL fibers were transfected with siRNAs against Huwe1 and were cultured for three days in 25mM or 2.5mM glucose media. Stem cell differentiation was accessed by Pax7/MyoG co-staining (n=3). Note that 1mM pyruvate was added in all experiments. Error bars indicated S.E.M. Paired Student’s t-test (two-tailed) was performed. *, p<0.05. **, p<0.01.

To test the effect of Huwe1 depletion in more physiologically relevant setting, we cultured muscle stem cells associated with muscle fibers. In this *ex vivo* system, muscle stem cells spontaneously activate and proliferate, forming colonies on top of the myofibers that contain undifferentiated and differentiating cells (Fig. 5D, left diagram). To avoid potential problems in stem cell activation, fibers were cultured in 25mM glucose during the first 24 hours, and subsequently transferred to either low or high glucose medium. To confirm that lowering the glucose concentration also impairs differentiation of the muscle stem cells in fiber culture, we determined the ratio of Pax7^+^ (undifferentiated) to MyoG^+^ (differentiated) cells in the colonies (Fig. EV4A). Differentiation was reduced in 5mM glucose and strongly impaired in 2.5mM glucose. Notably, low glucose did not affect the number of cells in the colonies, indicating that a shift to low glucose at 24 hours of culture did not impair cell proliferation (Fig. EV4B). Next, we tested whether siRNA-mediated depletion of Huwe1 restored differentiation in low glucose. We observed a significant increase in the proportion of MyoG-expressing cells in 2.5mM but not in 25mM glucose after siRNA treatment (Fig. 5D, right graph). In sum, Huwe1 mediates the effect of glucose on myogenic differentiation in both monolayer cultures and primary muscle stem cells associated with cultured myofibers; the latter is thought to model *in vivo* situation.

### PKM2-MyoD axis as a node controlling myogenic differentiation

Here we identified a new metabolic mechanism acting in myoblasts which links glucose availability to differentiation. Single-cell RNA-Sequencing of muscle stem cells showed that quiescent muscle stem cells express low level of mRNAs encoding glycolytic enzymes like PKM2 (Dell’Orso et al., 2019). Furthermore, MyoD protein is also absent in quiescent stem cells. As stem cells progress into activated state, MyoD protein needs to be precisely regulated in order to control the balance between self-renewal or terminal differentiation. We show here that PKM2 participates in such regulatory mechanisms. In addition to fructose 1,6-bisphosphate, the equilibrium of PKM2 tetramers/dimers is controlled by diverse intra- and extracellular cues such as growth factors, serine and ROS (Anastasiou et al., 2011, Hitosugi et al., 2009, Yang et al., 2012, Chaneton et al., 2012, Keller et al., 2012). Thus, the PKM2-MyoD axis might act as the nexus that integrates information about dynamic cellular environment and connect them to cell fate decision.

## Materials and Methods

### Cell culture and nutrient starvation

C2C12 and 293T cells (both from ATCC) were maintained in high glucose DMEM media (Thermofisher 11965092) supplemented with heat inactivated 10% FBS (Sigma F7524) and penicillin/streptomycin (Sigma P4333) at 37°C and 5% CO2. To induce differentiation, C2C12 cells were washed twice with PBS, and DMEM media containing 2% horse serum (Thermofisher 16050122) and penicillin/streptomycin was added. Differentiation media was replenished every two days. To modulate glucose concentration in the media, we used glucose and pyruvate free DMEM (Thermofisher 11966025), which was supplemented with indicated concentration of glucose (Thermofisher A2494001).

To induce acute energy stress, 25mM 2-deoxyglucose (Sigma D8375) was added to glucose and pyruvate free DMEM media. For differentiation assays, glucose and pyruvate free media was further supplemented with 1mM pyruvate (Thermofisher 11360070). To induce amino acid starvation, cells were washed twice with EBSS (Thermofisher 14155063) and incubated with EBSS buffer supplemented with 4.5g/L glucose, 1mM pyruvate and 10% FBS for one hour. FBS was dialyzed overnight against EBSS to deprive amino acid in the serum. For no starvation control, 50X amino acid mixture (Sigma M5550-100ML) was added.

### siRNA transfection

For siRNA transfection, we used 20μM siRNA duplexes using Lipofectamine RNAi max (Invitrogen) according to manufacturer’s guideline. siRNAs against PKM2 were purchased from Thermofisher (catalog number 4390771; assays numbers s71678 and s201789). Scrambled control and siRNAs against Upf1 and Huwe1 were synthesized from Eurofins with dTdT overhang on both ends. The targeting sequences were the followings:

Scrambled 5’-CGUACGCGGAAUACUUCG A-3’
Mouse Upf1 #1 5’-GCUGCCAUGAACAUCCCUAUU-3’
Mouse Upf1 #2 5’-GCAGCCAAUGUGGAGAAGAUA-3’
Mouse Huwe1 #1 5’-CCGCACUGUGUUAAACCAGAU-3’
Mouse Huwe1 #2 5’-GCACUGCUCAUCAAAGAUGUU-3’

### Constructs and DNA transfection

Human PKM2 wild-type construct was purchased from Addgene (44242) and sub-cloned to pCMV-Tag2B plasmid (Agilent Technologies). PKM2 S437Y mutant was generated by site-directed mutagenesis and sub-cloned to the same plasmid. Mouse MyoD was cloned from E13.5 whole mouse embryo cDNA and confirmed by sequencing. Afterwards, MyoD was sub-cloned to pCMV-Tag2B plasmid. Plasmid transfection was performed using Fugene HD (Promega E2311) according to the manufacture’s guideline.

### Generation of stable cell lines

Generation of Ampkα1/2 dKO cells: Plasmids containing Cas9, GFP and gRNAs against mouse Ampkα1 (79004) and Ampkα2 (79005) were obtained from Addgene. C2C12 cells were first transfected with Ampkα1 gRNA and sorted with GFP. After culturing the sorted cells for 2 weeks, cells were transfected with Ampkα2 gRNA and GFP sorted. These cells were diluted to single cell and seeded into 96-well plates. Ampkα1/2 dKO cells were screened by Western blotting.

Huwe1 stable knockdown: The same targeting sequences as for siRNA studies were cloned to pLKO.1 puro plasmid (Addgene 8453). As control, pLKO.1 puro plasmid containing scrambled sequence was used (Sigma SHC016-1EA). pLKO.1 constructs were co-transfected with psPAX2 (Addgene 12260) and VsvG (Addgene 8454) in 4:3:1 ratio to 293T cells. We changed the media 6 hours after transfection and collected the media for the next 48 hours. Supernatants were cleared by centrifugation at 1,800rcf for 10 minutes. Target cells were transduced with the supernatant containing 5μg/ml polybrene (Millipore TR-1003-G). Selection was started 24 hours after transduction with 3μg/μl puromycin (Sigma P8833).

### Streptolysin-O experiment

Streptolysin-O (SLO) was purchased from Sigma (S5265). SLO was reconstituted in 1ml pure water (Sigma W3513), aliquoted and stored in −20°C until future use. Cells were washed twice with 1X HBSS (Thermofisher 14175-053) and HBSS containing SLO (50X) was added. Cells were incubated in the incubator for 2-3 minutes. The optimal time was determined by checking one dish under the microscope after SLO treatment. After removing SLO, cells were gently washed twice with PBS and incubated with indicated media for 2 hours. Glycolytic intermediates were purchased from Sigma: Fructose 1,6-bisphosphate (F6803), DHAP (D7137) and G3P (39705).

### Cell lysis, immunoprecipitation, ubiquitination assay and Western blotting

Harvested cells were resuspended in lysis buffer (50mM Tris-Cl pH 7.5, 150mM NaCl, 1mM MgCl2, 1% NP40) supplemented with protease inhibitor (Roche 11836170001) and phosphatase inhibitor cocktail (Sigma P5726) and incubated on ice for 20 minutes. Lysates were centrifuged at 25,000rcf for 20 minutes on 4°C, and supernatants were transferred to new tubes. Protein concentration was measure by Bradford method (Bio-Rad 5000006) and boiled in Laemmli buffer for 10 minutes.

For immunoprecipitation, 2mg of cleared lysates were incubated with 3μg anti-Flag antibody (Sigma F3165) for 2 hours at 4°C with constant rotation. 20μl Dynabeads Protein G (Invitrogen 10004D) were added and incubated for another hour at 4°C. Beads were washed four times with lysis buffer containing different NaCl concentrations (150mM, 300mM, 500mM and 150mM) using magnetic stand. After removing last washing buffer, beads were boiled.

For ubiquitination assay, cells were treated with 20μM MG132 (Sigma M8699) for 4 hours before harvesting. Cells were lysed in RIPA buffer (50mM Tris-Cl pH 7.5, 150mM NaCl, 1mM MgCl2, 1% SDS and 0.1% Sodium Deoxycholate). Concentration was measured by BCA method (Thermofisher 23225). 3mg of cleared lysates were incubated with 3μg antibody against MyoD (Santa cruz sc-32758) or mouse IgG (Cell Signaling 5415) overnight at 4°C. After bead incubation, beads were washed 4 times with RIPA buffer and boiled.

Boiled samples were fractionated on SDS-PAGE gel, transferred to 0.45μm pore nitrocellulose membranes (GE healthcare), blocked for one hour with 5% skim milk (Sigma 70166) in PBST and overnight incubated with primary antibodies diluted in 5% BSA (Sigma A2153) in filtered PBST. After three times washing with PBST, membranes were treated with HRP-conjugated secondary antibodies (Cell Signaling 7074 and 7076) diluted at 1:5000 in skim milk for one hour at RT. After three times washing, membranes were developed using ECL primer solution (GE healthcare RPN2236). The primary antibodies used in this study are the followings: Muscle Creatine Kinase (Abcam ab54637, 1:2000), Troponin-T (Sigma T6277, 1:1000), MyoG (Santa Cruz sc-12732, 1:500), MyoD (Santa cruz sc-32758, 1:250), β-actin (Cell Signaling 4970, 1:1000), S6K (Cell Signaling 2708, 1:1000), phospho-S6K (Cell Signaling 97596, 1:1000), Ubiquitin (Abcam ab7780, 1:2000), AMPKα (Cell Signaling 2532, 1:1000), phospho-AMPKα (Cell Signaling 2531, 1:500), Lamin B (Abcam ab16048, 1:2000), β-Tubulin (Cell Signaling 2128, 1:2000), Huwe1 (Abcam ab70161, 1:1000) and PKM2 (Cell Signaling 4053, 1:1000).

### Reverse transcription and quantitative PCR

Harvested cell pellets were resuspended in 1ml Trizol reagent (Invitrogen) and RNA was isolated according to manufacturer’s guideline. 1μg RNA was reverse transcribed using random hexamer primer (Thermofisher N8080127) and Protoscript Reverse Transcriptase enzyme (NEB M0368). cDNA was dilute in water in 1:5 ratio. 1μl sample was used for one quantitative PCR reaction using 2X Cyber green mix (Thermofisher) and CFX96 (Biorad). The qPCR primers used in this study are the following:

β-actin For: 5’-GGCTGTATTCCCCTCCATCG-3’
β-actin Rev: 5’-CCAGTTGGTAACAATGCCATGT-3’
MyoD For: 5’-CCACTCCGGGACATAGACTTG-3’
MyoD Rev: 5’-AAAAGCGCAGGTCTGGTGAG-3’
MyoG For: 5’-GAGACATCCCCCTATTTCTACCA-3’
MyoG Rev: 5’-GCTCAGTCCGCTCATAGCC-3’

### Nucleocytoplasmic fractionation

Harvested cells were resuspended in hypotonic buffer (20mM HEPES, pH 7.9, 2mM KCl and 1mM DTT) and incubated on ice for 30 minutes. Triton X-100 was added to final concentration of 0.3% and briefly vortexed for 10 seconds. After one-minute centrifugation at 25,000rcf on 4°C, supernatants (cytosolic fractions) were transferred to new tubes. The pellets were washed once with hypotonic buffer and once with PBS. Hypertonic buffer (5mM HEPES, pH 7.9, 1.5mM MgCl2, 500mM NaCl, 0.2mM EDTA and 0.5mM DTT) was then added, and pellets were vortexed every five minutes at least 7-8 times. Pellets were centrifuged at 25,000rcf for 20 minutes on 4°C, and supernatants (nuclear fractionation) were transferred to new tubes.

### Immunostaining

Cells were cultured on coverslips coated with gelatin (Sigma G1393). After aspirating the media, cells were washed once with PBS and fixed with 4% PFA for 15 minutes on room temperature (RT). Fixed cells were permeablized with 0.3% Triton X-100 for 10 minutes on RT and washed once with PBS. Cells were blocked with 3% BSA in PBS for one hour on RT. Primary antibodies were diluted in blocking buffer, added on coverslips in humid chamber, and incubated overnight in cold room. After three times washing with PBS (10 minutes each), Cy2- or Cy3-conjugated secondary antibodies and DAPI (Sigma D9542) were diluted in blocking buffer and treated on the cells for one hour on RT. Samples were washed three times and mounted (Thermofisher 9990412). Primary antibodies used in this study are anti-Huwe1 (Bethyl A300-486A, 1:500) and MyoG (same as Western blot, 1:200). Secondary antibodies were from Jackson Immunoresearch (1:500).

### Myofiber culture and immunostaining

Myofibers from extensor digitorum longus (EDL) muscle were isolated as described before (Vogler et al., 2016). Briefly, dissected EDL muscles were digested with 0.2% Collagenase I (Sigma C0130) dissolved in high glucose DMEM media for 90 minutes at 37°C with gentle shaking. Digested fibers were transferred to high glucose DMEM media (without Collagenase I) and incubated for 30 minutes at 37°C. Single fibers were liberated by gently triturating with large diameter pastier pipet. Isolated fibers were transferred to high glucose DMEM supplemented with 10% heat inactivated horse serum, 1% chick embryo extract (MP bio MP 2850145) and Penicillin/Streptomycin. One hour after isolation, fibers were transfected with siRNAs against control or Huwe1. 24 hours later, media were removed, washed once with PBS, and replaced to the same media but with different glucose concentration. Fibers were cultured for another 72 hours until fixation. Immunostaining of myofibers were carried out using the same procedure as cell immunostaining except the following modifications; washing was performed with PBS plus 0.025% Tween-20, and HS/BSA solution was used for blocking (3% horse serum, 0.25% BSA and 0.1% Triton X-100 in PBS). The primary antibodies used for this experiment were MyoG (same as for cell immunostaining, 1:200) and Pax7 (homemade, 1:50). Pax7 antibody was previously described (Muller et al., 2002).

### Acquisition of fluorescence images

Fluorescence was visualized by laser-scanning microscopy (LSM700, Carl-Zeiss) using Zen 2009 software. Images were processed using ImageJ and Adobe Photoshop and assembled using Adobe Illustrator.

### Other chemicals

Other chemicals or reagents used in this study were Cyclohexmidie (Sigma 01810), EX527 (Selleckchem S1541), MLN4924 (Selleckchem HY-00062), DASA-58 (Selleckchem S7928) and TEPP-46 (Axon Medchem 2240).

## Acknowledgement

We thank Bettina Brandt for her technical assistance. M.K. was supported by post-doctoral fellowships from the Alexander von Humboldt Foundation and AFM-Telethon. This work was funded by Helmholtz society (to C.B).

## Author contributions

M.K conceived the project, conducted the experiment, and analyzed the data. Y.Z. performed the experiment in Fig. EV4. M.K and C.B wrote the manuscript.

## Declaration of interest

The authors do not have any conflicting interests.

---

**Figure Extended View 1.**
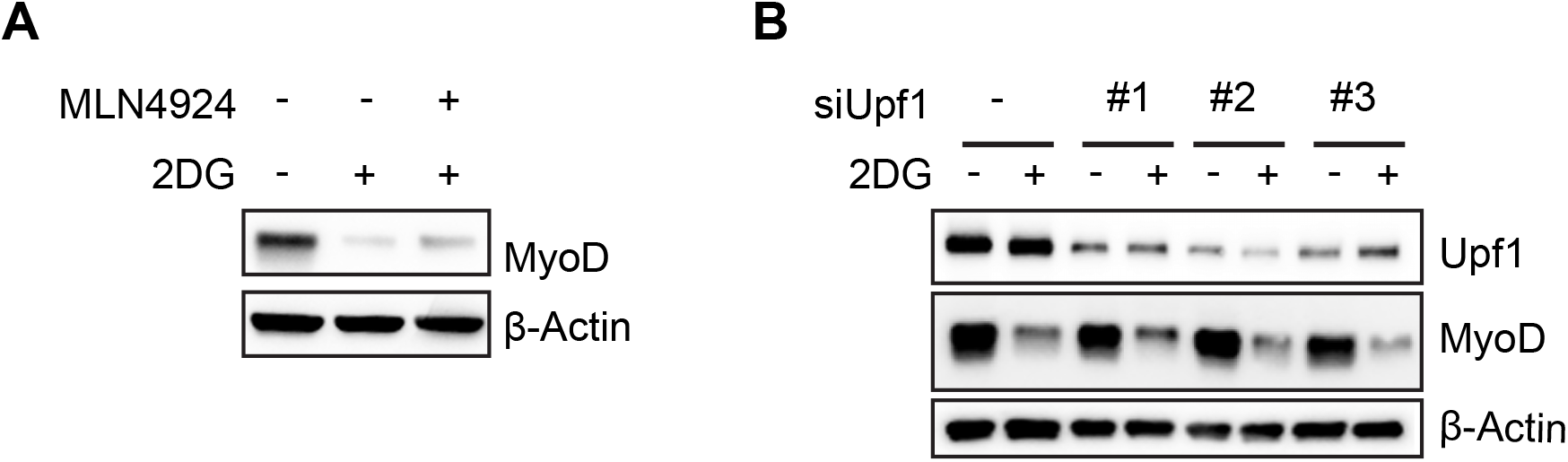
Atrogin-1 and Upf1 are not involved in the regulation of MyoD by 2DG. (A) Cells were co-treated with 2DG and 10μM MLN4924 (Atrogin-1 inhibitor) for 4 hours. (B) Cells were transfected with two different siRNAs against Upf1 and treated with 2DG for 4 hours. All results were validated in two or more independent experiments.

**Figure Extended View 2.**
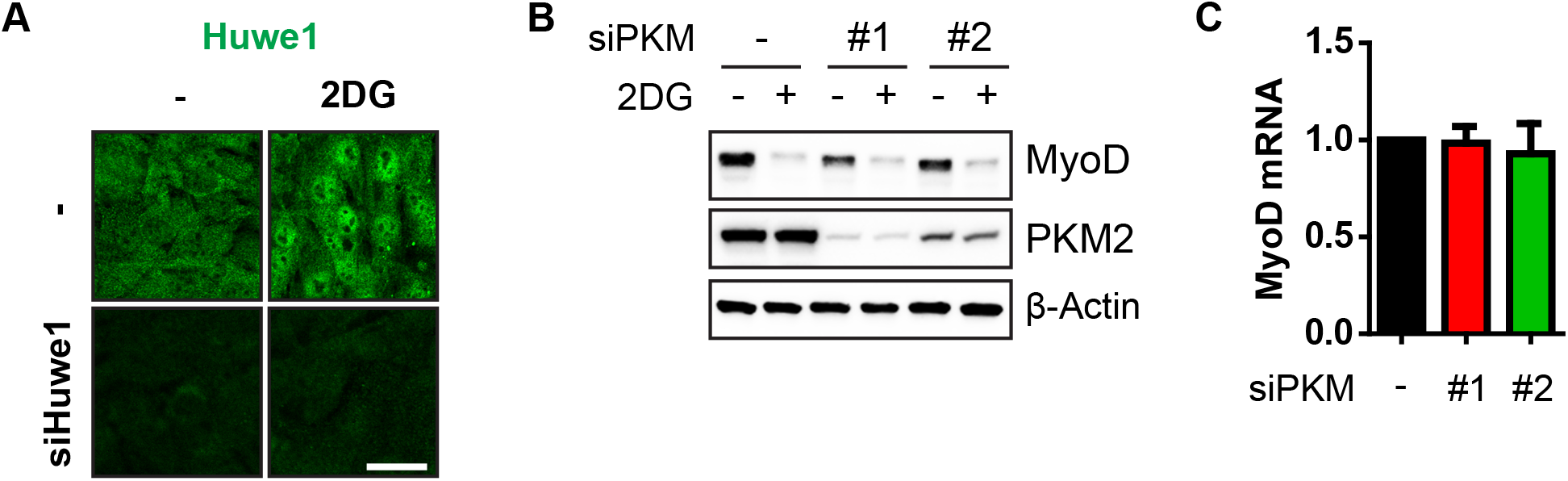
Loss of PKM2 decreases MyoD protein. (A) Huwe1 immunostaining in Huwe1 siRNA transfected cells validate the specificity of the antibody. Scale bar, 20μm. (B) C2C12 cells were transfected with two different siRNAs against PKM2 and treated with 2DG for 4 hours. (C) Samples from (B) were analyzed for MyoD mRNA level by RT-qPCR (n=3).

**Figure Extended View 3.**
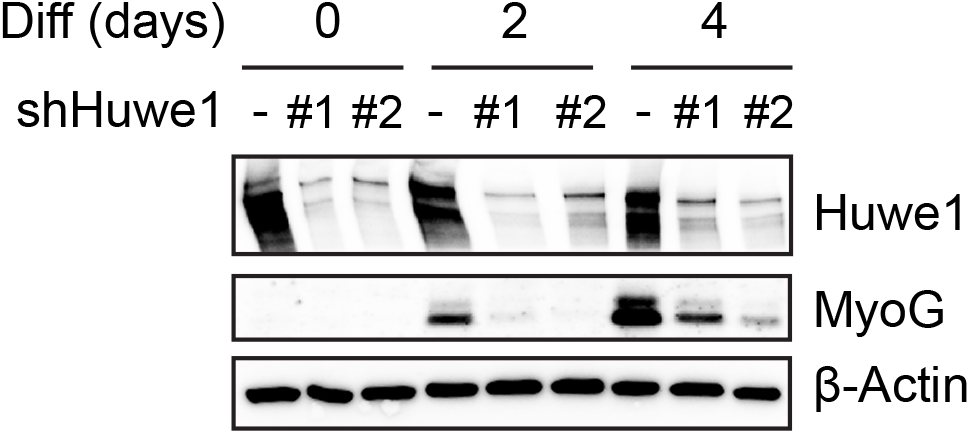
Prolonged loss of Huwe1 can impair myogenic differentiation. C2C12 cells with stable knock-down of Huwe1 by shRNA (maintained more than 2 weeks) were induced to differentiate in 25mM glucose media condition.

**Figure Extended View 4.**
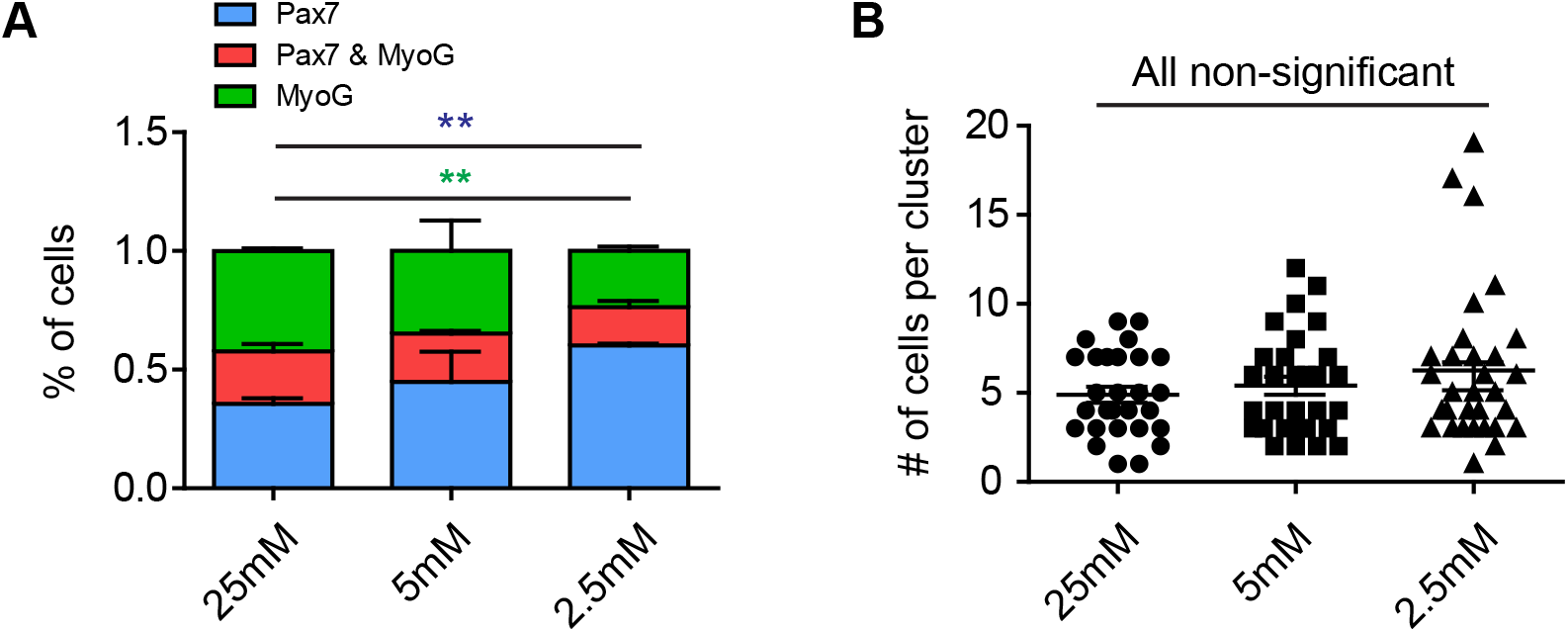
Lowering glucose impairs differentiation in cultured EDL myofibers. (A) Isolated mouse EDL fibers were first cultured in 25mM glucose media for 24 hours. On the second day, media were switched to 25mM, 5mM, or 2.5mM glucose and cultured for three more days. Differentiating and undifferentiated stem cells were detected by immunostaining for MyoG and Pax7, respectively (n=3). (B) This culture condition does not affect cell proliferation. Numbers of cells per myogenic colony are quantified (n=3). Note that 1mM pyruvate was added in all experiments. Error bars indicated S.E.M. Paired Student’s t-test (two-tailed) was performed. *, p<0.05. **, p<0.01.

## Notes

### Competing Interest Statement

The authors have declared no competing interest.

